# Novel Requirement for Staphylococcal Cell Wall-Anchored Protein SasD in Pulmonary Infection

**DOI:** 10.1101/2022.04.01.486802

**Authors:** Jennifer A Grousd, Abigail M. Riesmeyer, Vaughn S. Cooper, Jennifer M. Bomberger, Anthony R. Richardson, John F. Alcorn

**Author notes:** Address correspondence to Dr. John F. Alcorn.

## Abstract

*Staphylococcus aureus* can complicate preceding viral infections, including influenza virus. A bacterial infection combined with a preceeding viral infection, known as super-infection, leads to worse outcomes compared to single infection. Most of the super-infection literature focuses on the changes in immune responses to bacteria between homeostatic and virally infected lungs. However, it is unclear how much of an influence bacterial virulence factors have in super-infection. Staphylococcal species express a broad range of cell wall-anchored proteins (CWAs) that have roles in host adhesion, nutrient acquisition, and immune evasion. We screened the importance of these CWAs using mutants lacking individual CWAs *in vivo* in both bacterial pneumonia and influenza super-infection. In bacterial pneumonia, lacking individual CWAs led to varying decreases in bacterial burden, lung damage, and immune infiltration into the lung. However, the presence of a preceding influenza infection partially abrogated the requirement for CWAs. In the screen, we found that the uncharacterized CWA *S. aureus* surface protein D (SasD) induced changes in both inflammatory and homeostatic lung markers. We further characterized a SasD mutant (sasD A50.1) in the context of pneumonia. Mice infected with sasD A50.1 had decreased bacterial burden, inflammatory responses, and mortalty compared to wildtype *S. aureus*. Mice also had reduced levels of IL-1β compared with wildtype, likely derived from macrophages. Reductions in IL-1β transcript levels as well as increased macrophage viability implicate altered macrophage cell death pathways. These data identify a novel virulence factor for *S. aureus* that influences inflammatory signaling within the lung.

**Importance:** *Staphylococcus aureus* is a common commensal bacteria that can cause severe infections, such as pneumonia. In the lung, viral infections increase the risk of staphylococcal pneumonia, leading to combined infections known as super-infections. The most common virus associated with *S. aureus* pneumonia is influenza, and super-infections lead to worse patient outcomes compared to either infection alone. While there is much known about how the immune system differs between healthy and virally infected lungs, the role of bacterial virulence factors in super-infection is less understood. The significance of our research is identifying new bacterial virulence factors that play a role in the initiation of infection and lung injury, which could lead to future therapies to prevent pulmonary single or super-infection with *S. aureus*.

## Introduction

Respiratory viral infections can be complicated by bacterial pneumonia, leading to increased rates of morbidity and mortality. *Staphylococcus aureus* has been shown to complicate several viral infections such as influenza, respiratory syncytial virus, and rhinovirus(1). This is also seen in the current viral pandemic with COVID-19, with one study finding a mortality rate over 60% for those infected with both SARS-CoV-2 and *S. aureus*(2). While *S. aureus* is considered to be a common commensal, it can cause severe disease such as endocarditis, bacteremia, sepsis, and death(3). Combined with a pre-existing viral infection (colloquially referred to as super-infection), it is unsurprising that outcomes in patients that are super-infected with *S. aureus* are worse than either bacterial or viral infection alone. The virus most commonly associated with *S. aureus* is influenza. Influenza is a seasonal respiratory virus that causes an estimated 294,000-518,000 deaths worldwide each year(4). Since 2009, *S. aureus* is the primary contributor to influenza bacterial super-infections(5–8).

Preceeding viral infections increase the susceptibility to *S. aureus* infection through three main mechanisms: 1) modifying the expression of host proteins involved in *S. aureus* adhesion or internalization, 2) synergism of viral- and bacterial-induced epithelial invasion and damage, and 3) reduced clearance of the bacteria by altering the immune response(1, 9–13). A large body of literature exists focused on the immunological differences in antibacterial immunity that occur with a preceding viral infection(12–14). In general, the antiviral response inhibits the clearance of a bacterial infection through various mechanisms(9–13). There are also physiological differences that occur due to viral infection that may increase susceptibility to secondary infections. Many of the bacteria known to cause super-infection are nasal commensals, and inflammation from influenza has been shown to increase dissemination from the nasopharynx to the lung(10, 15). The virus itself, and the immune response to the virus, can lead to epithelial and endothelial damage, which could lead to increased nutrient resources as well as potentially expose cryptic receptors for bacterial adherence to cells or basement membrane components(9, 10, 16). Influenza neuraminidase and wound healing responses can alter the cellular expression of receptors on cells, which may act as adhesion sites for bacteria(9, 10) (17, 18). Once the bacteria are attached, bacterial toxins could synergize with the virus to cause further damage and inflammation in the lung, potentially leading to the increased morbidity and death in super-infection(9, 10, 19).

Few studies have focused on the bacterial side of super-infections. For *S. aureus* specifically, the role of few virulence factors have been described within the lung, even within the context of *S. aureus* pneumonia. Because the super-infection literature suggests that viral infection can influence bacterial adherence, we explored surface proteins of *S. aureus*, collectively known as the cell wall-anchored proteins (CWAs), since many of these proteins have known roles in adhesion(20, 21). *S. aureus* can express up to 24 CWAs, with the most prevalent subfamily known as microbial surface component recognizing adhesive matrix molecule (MSCRAMM) proteins(20, 21). CWAs are covalently attached to the cell wall via the Sortase A enzyme that recognizes the cell wall sorting motif LPXTG(20, 22–24). Because these are surface exposed proteins, they are in contact with the host and have a variety of known functions such as host adhesion, biofilm formation, immune evasion, and nutrient acquisition for both colonization and invasive infection(20, 21). Some CWAs have been described to play a part in nasal colonization(20, 21, 25). Since these CWA proteins play an important part in *S. aureus* colonization and infection, we decided to screen several CWA members in the lung in both bacterial pneumonia and influenza super-infection.

## Results

### Screening Cell Wall-Anchored Protein Mutants During Bacterial Pneumonia and Influenza Super-infection

Since cell wall-anchored proteins (CWAs) are exposed on the cell surface of *S. aureus*, we hypothesized that these proteins may be playing a role in colonization and/or infection in the lung. We screened nine CWA mutants (*fnbB::Tn, clfA::Tn, clfB::Tn, sdrC::Tn, sdrD::Tn, sdrE::Tn, isdB::Tn, sasG::Tn*, and *sasD::Tn*) and the corresponding wildtype (WT) strain JE2 in the context of bacterial pneumonia and influenza super-infection. We also included a Sortase A (*srtA::Tn*) mutant, the enzyme responsible for attaching these CWAs to the cell wall of *S. aureus*. In terms of bacterial burden within the lung (Figure 1 A), lacking individual CWA proteins during bacterial pneumonia lead to varying decreases in burden when compared to the WT strain. The differences in bacterial burden did not appear to be due to differences in *in vitro* growth rates of the various mutants (Supplementary Figure 1 A and B). Interestingly, the Sortase A mutant did not have a significant decrease in bacterial burden. The mutant lacking SasG (*S. aureus* surface protein G; *sasG::Tn*) had the largest decrease in burden during bacterial pneumonia. During super-infection, mutants had increased burden versus single infection, although only the WT and ClfA mutant (clumping factor A; *clfA::Tn*) were significantly increased. The mutant lacking IsdB (iron-regulated surface determinant protein B; *isdB::Tn*) was the only mutant that had significantly decreased burden in both bacterial pneumonia and super-infection. Lacking *S. aureus* CWAs did not impact viral burden in the lung (Supplementary Figure 1 C).

**Figure 1:**
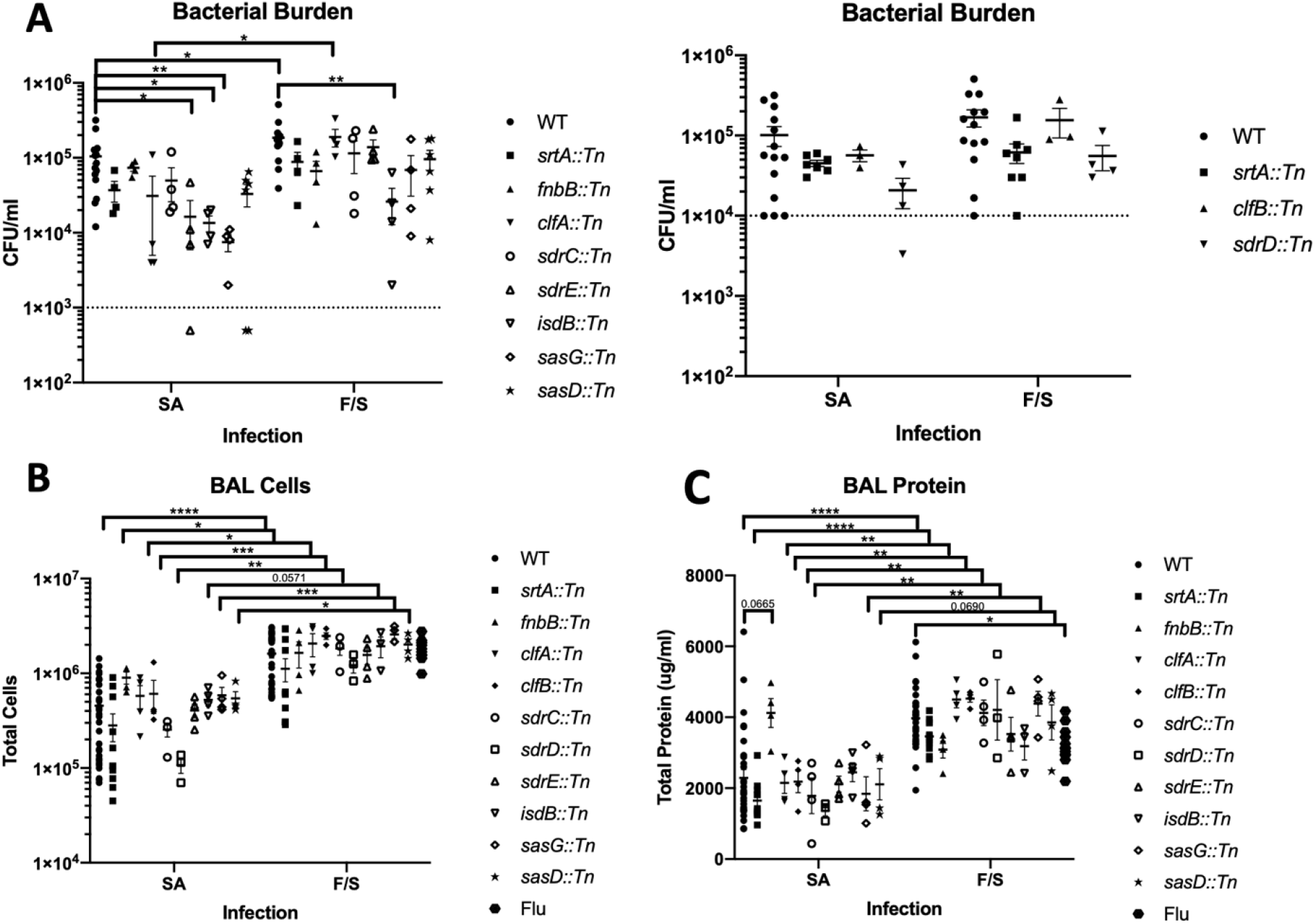
Differential Impact of *S. aureus* CWA Mutants in Bacterial Pneumonia and Influenza Super-Infection. Mice were inoculated with PBS or 100 PFU of influenza on day 0 and six days later were infected with PBS or 1×10 CFU of WT MRSA or a strain lacking individual CWA (see graphs) and harvested 24 hours later. **A.** Bacterial burden in bacterial pneumonia (SA) or influenza super-infection (FS) 24 hours post MRSA infection. Mice with undetectable CFU were graphed as half of the limit of detection. **B-C.** Total cells (**B**) or total protein (**C**) in the bronchoalveolar lavage (BAL) in bacterial pneumonia (SA) or influenza super-infection (FS). Statistics tested by Two-way ANOVA with Sidak’s multiple comparison correction. * p<0.05, ** p<0.01, ***p<0.001, ****p<0.0001. N=2-4, combination of several experiments, data graphed as mean ± SEM (standard error of the mean). srtA: Sortase A, fnbB: fibronectin binding protein B, clfA/B: clumping factor A/B, sdrC/D/E: serine-aspartate repeat containing protein C/D/E, isdB: iron-regulated surface determinant B, sasD/G: *S. aureus* surface protein D/G.

To look at lung inflammation with CWA mutants, we examined the number of cells in the airway via bronchoalveolar lavage (BAL) (Figure 1 B). Cellular immune infiltrates in the lung varied based on the mutant. During super-infection, almost all mutants had significantly higher numbers of BAL cells in the airways. The number of cells in the BAL during super-infection was very similar to the influenza alone level at day 7 post infection. To look at acute lung injury and leak, we measured total protein in the BAL (Figure 1 C). Although not as variable, the level of protein in the BAL during bacterial pneumonia varied based on the mutant. Interestingly, the mutant lacking FnbB (fibronectin binding protein B; *fnbB::Tn*) had the highest level of protein in the BAL in bacterial pneumonia. Lung leak also significantly increased during super-infection compared to bacterial pneumonia. Only the WT had significantly increased BAL protein during super-infection compared to influenza alone.

### Immune Responses to CWA Mutants in Bacterial Pneumonia and Influenza Super-Infection

Next, we examined the inflammatory response to CWA mutants via cytokines in lung homogenate. To determine if immune signatures were similar in CWA subfamilies, we visualized the cytokine data by clustering analyses (Figure 2 A and B). In bacterial pneumonia, there were three clusters of inflammatory responses which we termed: low inflammation, mixed inflammation, and high inflammation (Figure 2 A). Unsurprisingly, the Sortase A mutant, which lacks all CWAs on the cell surface, had the lowest level of cytokine induction. *srtA::Tn* had significant decreases in type 2 cytokines IL-9 (p<0.0001) and IL-13 (p=0.0025), but not IL-4 and IL-5. This was not driven by IL-33 expression, as *srtA::Tn* trended towards increased IL-33 expression via qPCR (p=0.0710) (data not shown). The other mutant in the low inflammation cluster was *sdrD::Tn* (serine aspartate repeat containing protein D), which had higher levels of expression of type 1 and type 2 cytokines compared to *srtA::Tn*. The mutants found in the mixed inflammation cluster typically had higher levels of innate immunity cytokines and chemokines, but lower levels of type 1, 2, and 17 cytokines. The high inflammation cluster, which contained the WT strain as well as most Clf and Sdr members, had the highest levels of cytokines. During super-infection, the clustering of CWAs by cytokine expression was very similar to bacterial pneumonia (Figure 2 B). Again, the strains cluster into three groups distinct from influenza alone and the only mutant that switched clusters was *sasG::Tn*.

**Figure 2:**
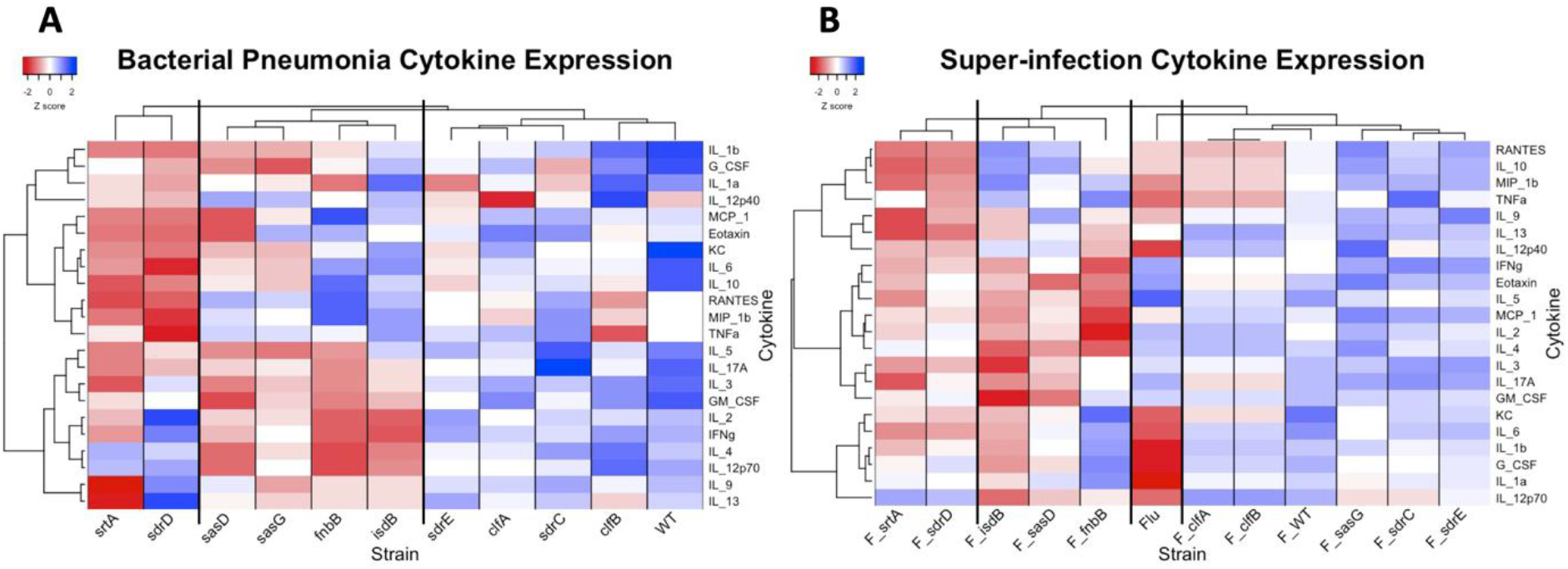
CWA Mutant Induced Cytokine Expression in Bacterial Pneumonia and Influenza Super-Infection. **A.** Cytokine expression in bacterial pneumonia. There are three clusters (from left to right): low inflammation, mixed inflammation, and high inflammation. **B.** Cytokine expression in super-infection. There are four clusters (from left to right): low inflammation, mixed inflammation, influenza alone (Flu), and high inflammation. Cytokines were measured via multiplex analyses of lung homogenate. For each cytokine, average values for each MRSA strain were log transformed and converted to Z scores. Heatmap clustering was performed by cytokine using the Pearson correlation and graphed using the heatmap.2 function of the gplots package in R. srtA: Sortase A, fnbB: fibronectin binding protein B, clfA/B: clumping factor A/B, sdrC/D/E: serine-aspartate repeat containing protein C/D/E, isdB: iron-regulated surface determinant B, sasD/G: *S. aureus* surface protein D/G.

### Characterization of SasD during Bacterial Pneumonia

We identified that the mutant lacking SasD (*S. aureus* surface protein D; *sasD::Tn*) induced reduced levels of cytokines G-CSF, CXCL1, MCP-1, and IL-1β compared to the WT strain during bacterial pneumonia (Supplemental Figure 2 A). Additionally, we also saw increased levels of epithelial and lung homeostasis marker gene expression including tight junction protein 1 (*tjp1*) and mucin 5b (*muc5b*) (Supplemental Figure 2 B). This suggested to us that this mutant may be causing less inflammation in the lung leading to improved epithelial and lung function. There is very little known about SasD and to our knowledge, there has been no *in vivo* characterization of this CWA. Therefore, we decided to characterize this mutant in the context of bacterial pneumonia. To eliminate the potential for unknown mutations in the transposon mutant, we transduced the mutant into the JE2 strain, creating the strain sasD A50.1. This strain had no difference in *in vitro* growth compared to the WT strain (Supplemental Figure 3 A and B). At 24 hours post infection, mice infected with sasD A50.1 had reduced bacterial burden and immune infiltrate in the BAL (Figure 3 A and B). Infection with sasD A50.1 led to a decrease and increase in percentage of neutrophils and eosinophils, respectively. While there were no changes in genes related to neutrophil function (Supplementary Figure 3 C), we did see a 50% reduction in the neutrophil:eosinophil ratio in the lung (Figure 3 C and D). The change in bacterial burden and immune cell infiltrate did not lead to differences in acute lung injury by BAL protein (Figure 3 E) or histological score (Supplemental Figure 3 D). However, we did see significant changes to transcripts related to lung homeostasis (Supplementary Figure 3 E). Additionally, we saw a significant delay in mortality in mice infected with sasD A50.1 compared to WT *S. aureus* (Figure 3 F). The changes seen 24 hours post infection in mice infected with sasD A50.1 may be because of a decrease in survival of the mutant in the lung, as this mutant had a reduced competitive index when mice were infected with a 1:1 ratio of mutant:WT bacteria (Figure 3 G).

**Figure 3:**
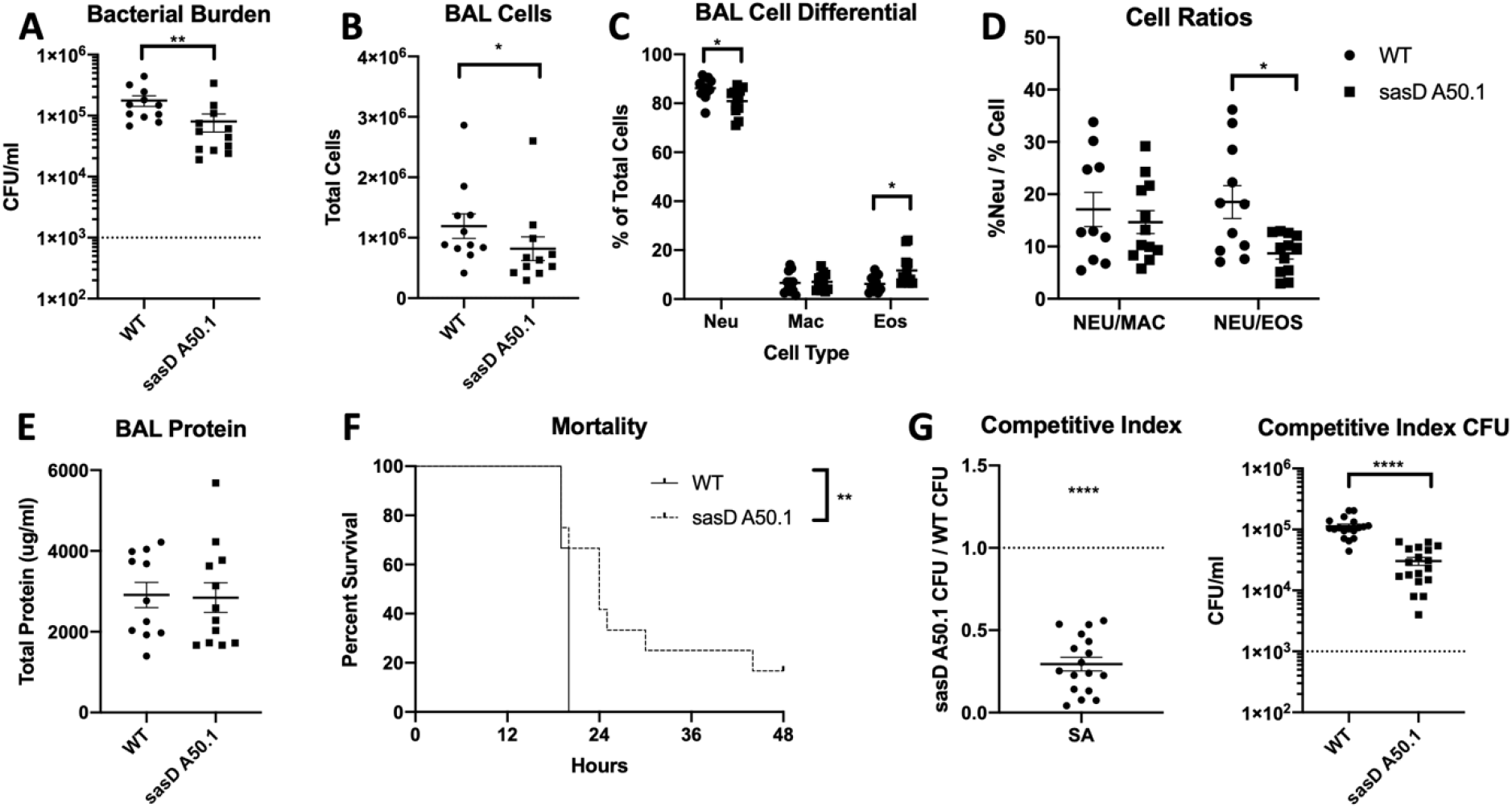
SasD is Required for *S. aureus* Bacterial Pneumonia. **A-E.** Mice were infected with 1×10^8^ CFU WT MRSA or MRSA lacking SasD (sasD A50.1) for 24 hours. **A.** Bacterial burden in mice infected with MRSA for 24 hours**. B.** Total cells in the bronchoalveolar lavage (BAL). **C-D.** Cell differentials (**C**) and neutrophil cell ratios (**D**) of BAL cells. **E.** Total protein in the BAL. **F.** Mice were infected with a lethal dose (2×10^8^ CFU) of WT or MRSA lacking SasD (sasD A50.1). **G.** Competitive index of WT and mutant sasD A50.1 MRSA in the lung. Mice were infected with a 1:1 ratio of WT:sasD A50.1 for a total dose of 1×10^8^ CFU for 24 hours. Whole lungs were collected in 2 ml PBS and homogenized and plated for CFU with and without antibiotic selection. Competitive index is calculated as the ratio of Mutant CFU:WT CFU at 24 hpi. Statistics tested by student Mann-Whitney (A,B, D, G), Two-way ANOVA with Sidak’s multiple comparisons (C), log-ranked Mantel Cox test (F), one sample T-test with H0 set to 1 (1:1 ratio of mutant:WT) (G). * p<0.05, ** p<0.01,**** p<0.0001. N=4-8, combination of several experiments, data graphed as mean ± SEM.

To evaluate the inflammatory state within the lung, we looked at cytokine expression in the lung 24 hours post infection. Similar to the original screen, we saw decreases in protein levels of IL-17A, CXCL1, GM-CSF, and IL-1β in mice infected with sasD A50.1 (Figure 4 A). We also saw a decrease in transcript levels for IL-17A, CXCL1, and IL-1β (Figure 4 B). Because we saw a decrease in IL-1β and *S. aureus* is known to induce the NLRP3 (NOD-, LRR- and pyrin domain-containing protein 3) inflammasome(26), we also looked at inflammasome components. We saw a significant decrease in NLRP3 transcript, but not the general inflammasome adaptor ASC (apoptosis-associated speck-like protein containing C-terminal caspase recruitment domain [CARD]; *pycard*) (Figure 4 B).

**Figure 4:**
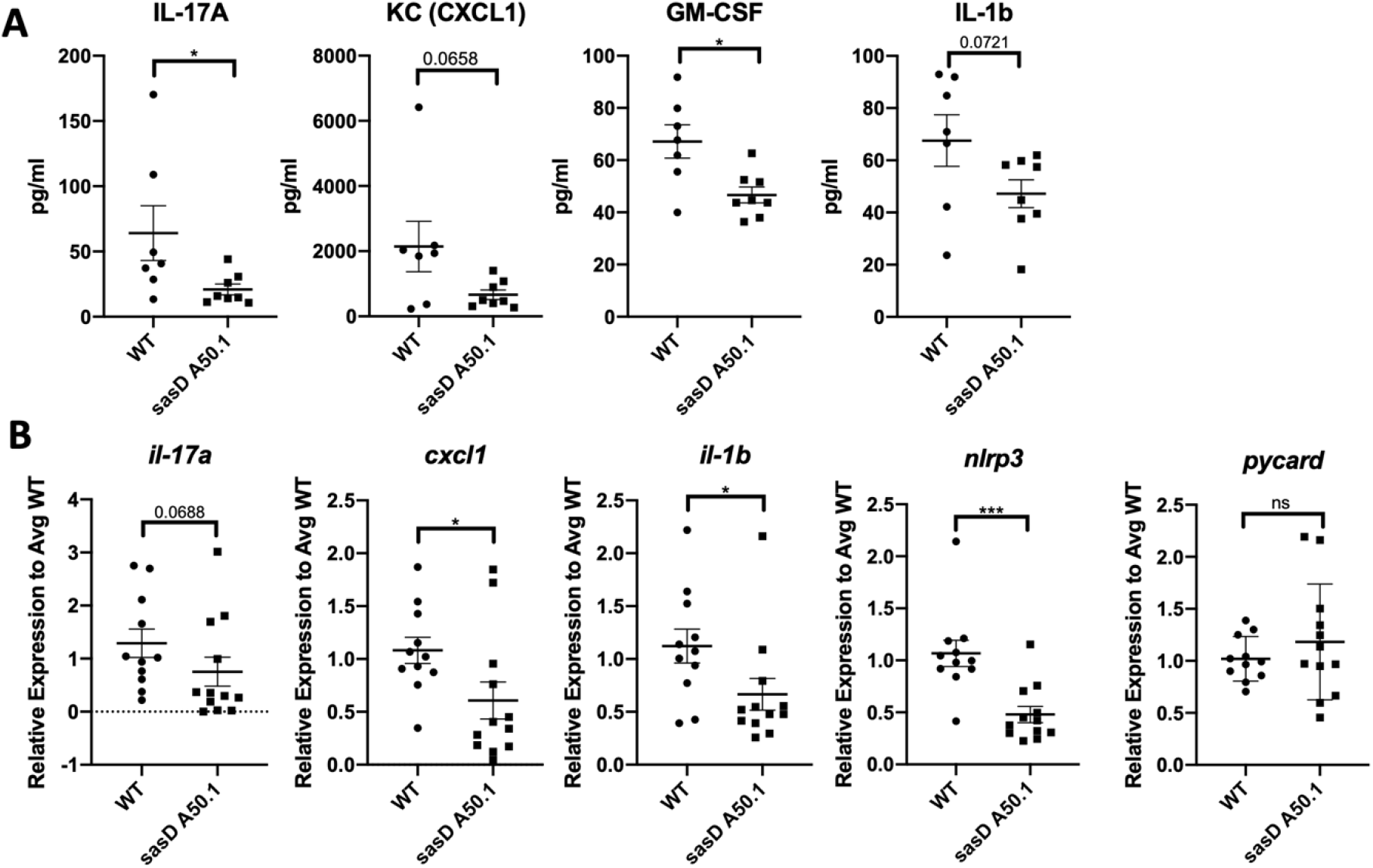
SasD is Required for Inflammation in Mice Infected with MRSA. Mice were infected with 1×10^8^ CFU WT MRSA or MRSA lacking SasD (sasD A50.1) for 24 hours. **A.** Cytokine protein levels in lung homogenate. **B.** Gene expression levels of cytokines and inflammasome components relative to average WT levels in the lung. Statistics done by Mann-Whitney test, * p<0.05, *** p<0.001. N=4, combination of several experiments, data graphed as mean ± SEM.

### Changes in Inflammation Occur Early During Infection With sasD A50.1

Mice infected with sasD A50.1 for 6 hours also had a reduction in bacterial burden compared with WT (Figure 5 A). While the total number of immune cells recruited into the airway was not different between WT and sasD A50.1 infected animals (Figure 5 B), we did see significant decreases in the percentages and total numbers of macrophages in the BAL (Figure 5 C-D). This increase in the neutrophil:macrophage ratio in the lung during this early timepoint (Figure 5 E) may be associated with less neutrophil recruitment later during infection. Unlike 24 hours post infection, we did see a significant difference in acute lung injury via protein in the BAL (Figure 5 F) and there was a significant difference in peribronchial inflammation via histology (Supplementary Figure 4 A). However, we did not see changes in lung homeostasis transcripts (Supplemental Figure 4 B). We also saw a survival defect of this mutant early on in infection via competitive index (Figure 5 G), which does not appear to be due to differences in early recruited neutrophil function (Supplemental Figure 4 C).

**Figure 5:**
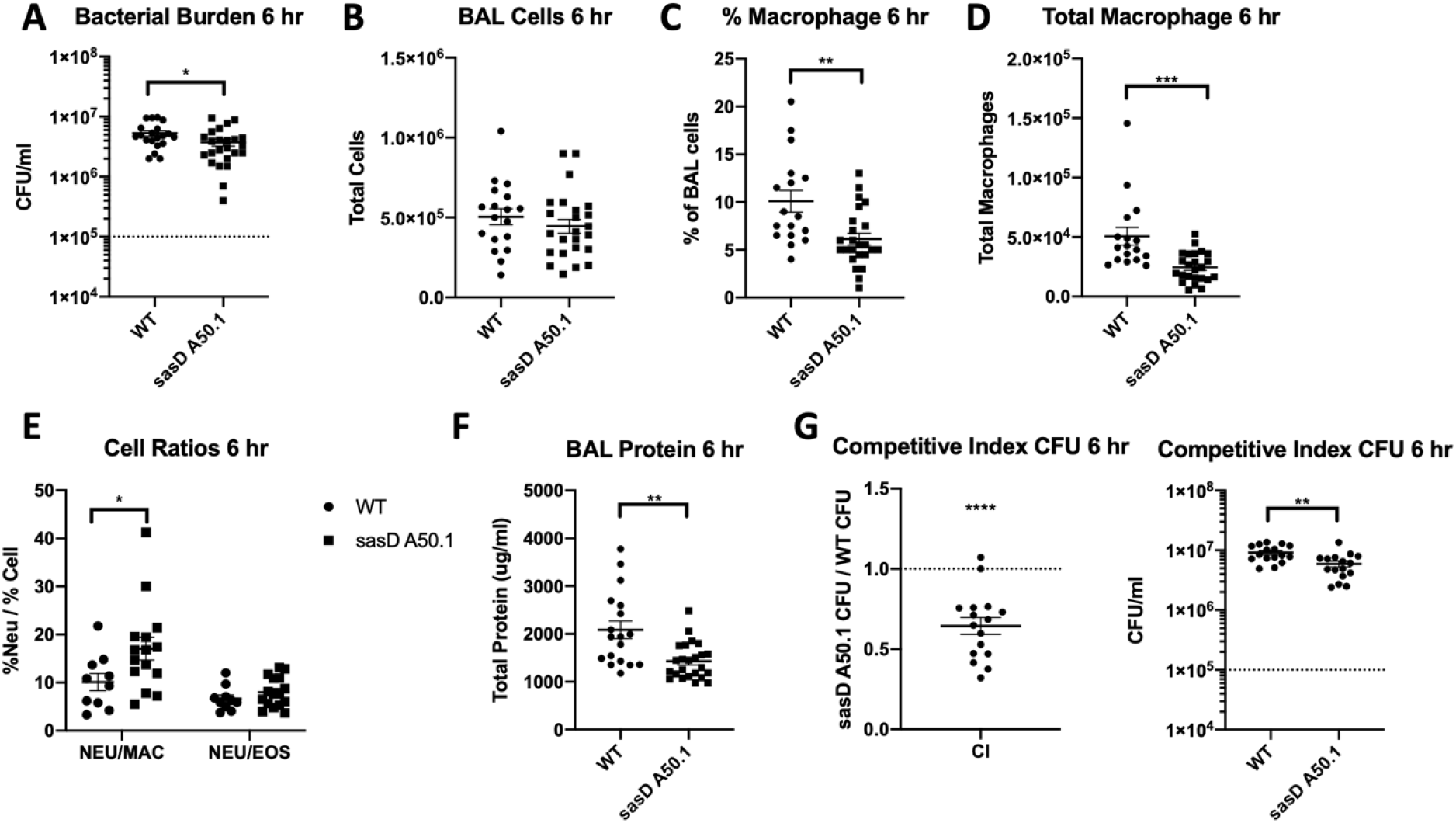
SasD Impacts Initiation of Host Defense against MRSA in Bacterial Pneumonia. **A-F.** Mice were infected with 1×10^8^ CFU WT MRSA or MRSA lacking SasD (sasD A50.1) for 6 hours. **A.** Bacterial burden in mice infected with MRSA for 6 hours. **B.** Total cells in the bronchoalveolar lavage. **C-D.** Percentage (**C**) and total number (**D**) of macrophages in the BAL. **E.** Neutrophil cell ratios in the BAL. **F.** Total protein in the BAL. **G.** Competitive index of WT and mutant sasD MRSA in the lung. Male and female mice were infected with a 1:1 ratio of WT:sasD A50.1 for a total dose of 1×10^8^ CFU for 6 hours. Whole lungs were collected in 2 ml PBS and homogenized and plated for CFU with and without antibiotic selection. Competitive index is calculated as the ratio of Mutant CFU:WT CFU at 6 hpi. Statistics tested by student Mann-Whitney (A, C, D, F, G), Two-way ANOVA with Sidak’s multiple comparisons (E), one sample T-test with H_0_ set to 1 (1:1 ratio of mutant:WT) (G). * p<0.05, ** p<0.01, ***p<0.001, **** p<0.0001. N=8, combination of several experiments, data graphed as mean ± SEM.

The inflammatory response to sasD A50.1 at 6 hours post infection was very similar to 24 hours post infection (Figure 6). Again, we saw reductions in IL-17A, and IL-1β with the addition of a decrease in G-CSF and IL-6 (Figure 6 A). By transcript levels, we saw reductions in both IL-17A and IL-23A (Figure 6 B), suggesting that the antibacterial immunity response is not as robust to sasD A50.1 compared to the WT at this time point, potentially due to the difference in survival between the two strains (Figure 5 G). We also saw a reduction in inflammasome components NLRP3 but not ASC (Figure 6 B).

**Figure 6:**
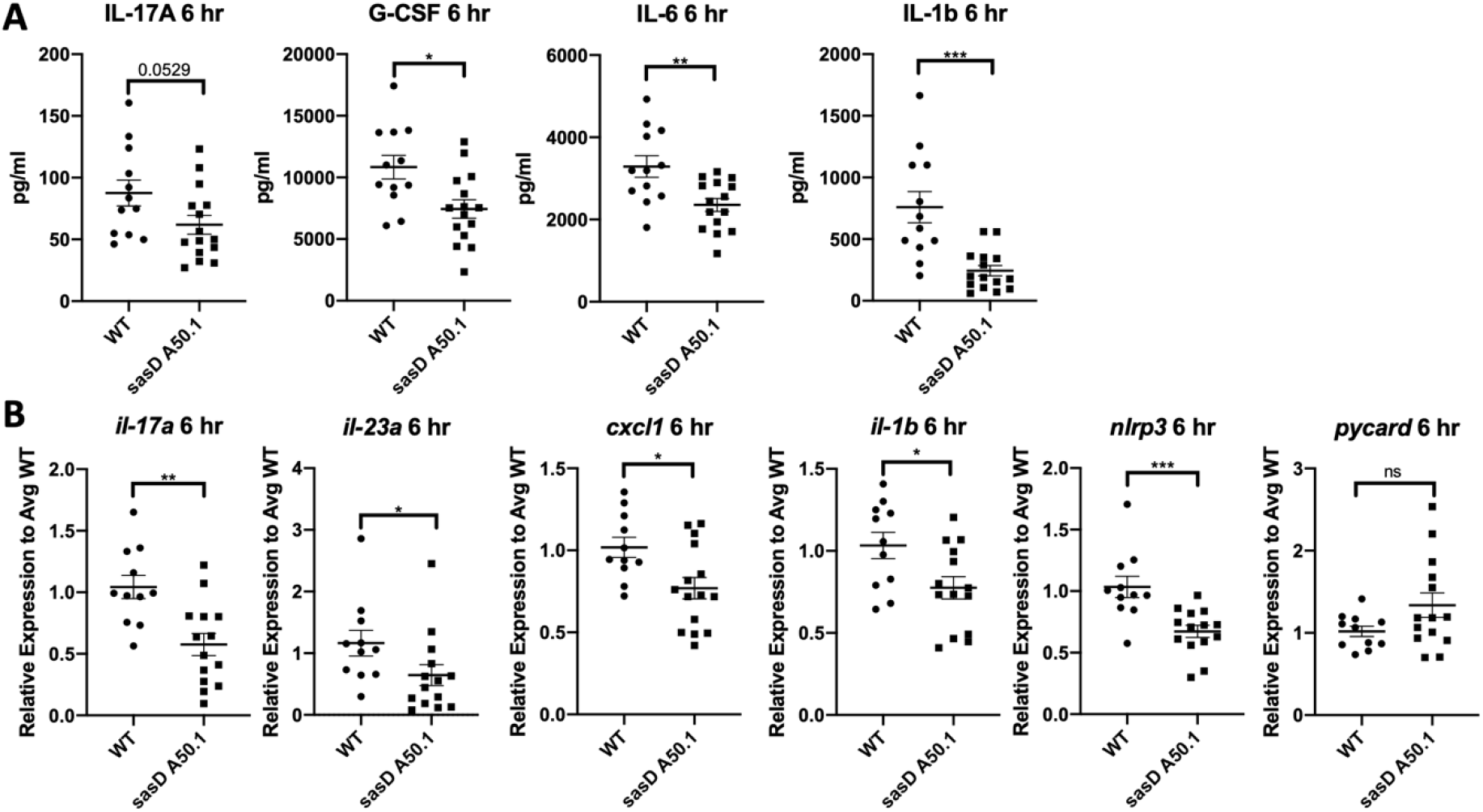
SasD is Required for Early Inflammation During Infection with MRSA. Mice were infected with 1×10^8^ CFU WT MRSA or MRSA lacking SasD (sasD A50.1) for 6 hours. **A.** Cytokine protein levels in lung homogenate. **B.** Gene expression levels of cytokines and inflammasome components relative to average WT levels. Statistics done by student Mann-Whitney, * p<0.05,**p<0.01, *** p<0.001. N=8, combination of several experiments, data graphed as mean ± SEM.

### Early Macrophage Interactions with sasD A50.1

Because of the differences seen in macrophages early on during infection, we looked at early macrophage interactions with sasD A50.1. Using the macrophage cell line RAW264.7, we saw a significant reduction in transcript expression of IL-1β and TNFα despite no difference in bacterial burden or amount of phagocytosed bacteria (Figure 7 A and B). In bone marrow-derived macrophages (BMDMs), infection with sasD A50.1 lead to an increase in macrophage viability compared to WT infected BMDMs at 3 hours post infection (Figure 7 C). At this timepoint we did not see any differences in pro-IL-1β, caspase 1, or caspase 1 cleavage (Figure 7 D and E).

**Figure 7:**
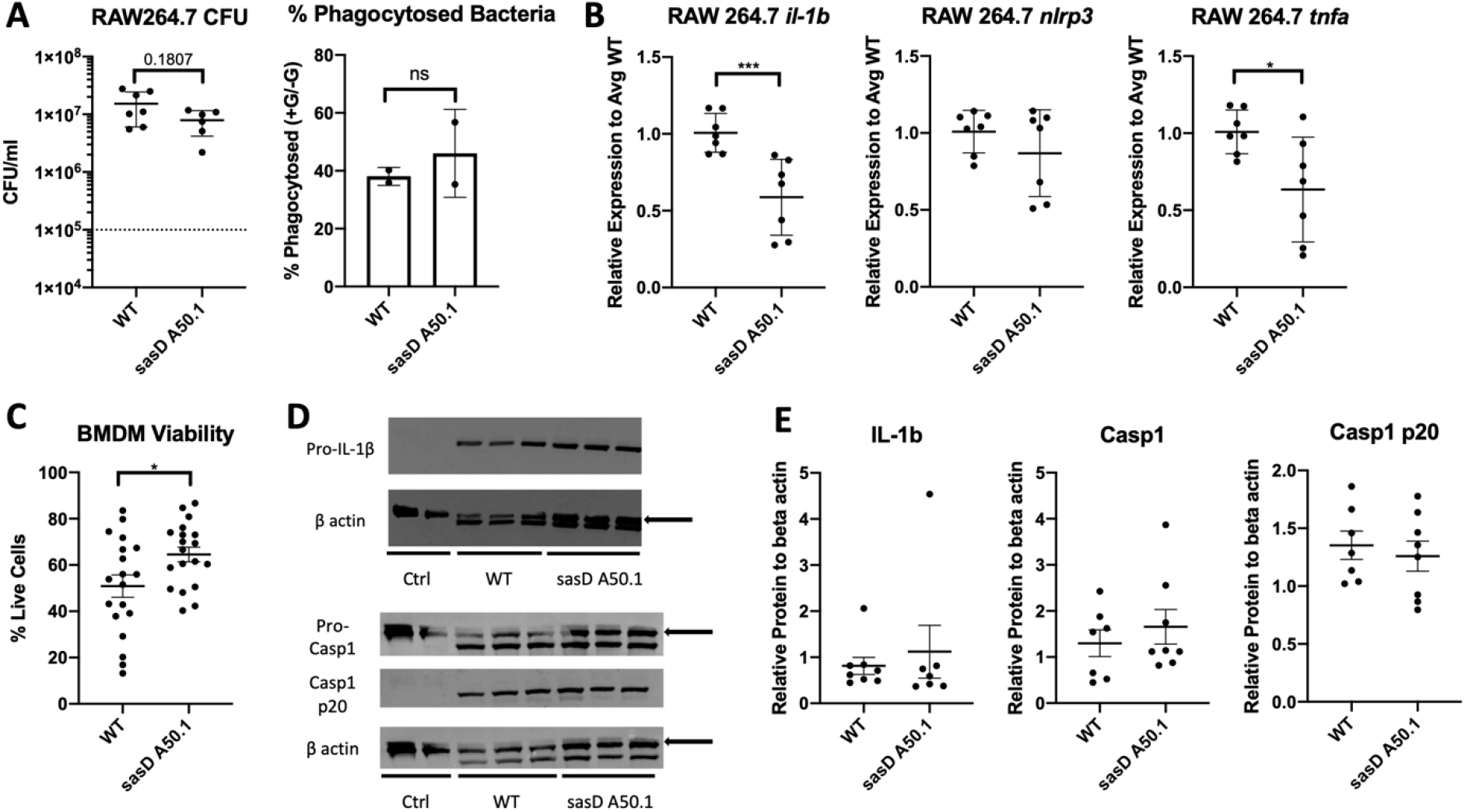
SasD Increases Macrophage Inflammation and Decreases Survival. **A-B.** RAW264.7 macrophages infected with WT or sasD A50.1 MRSA for 3 hours at an MOI of 10. Macrophages were infected for one hour in the absence of antibiotics, media was then replaced with antibiotic- and serum-free media with or without gentamicin for 1 hour, and changed to antibiotic free media. CFU and transcript graphs show without gentamicin conditions. **A.** Bacterial burden and % phagocytosed bacteria in RAW264.7 macrophages. % Phagocytosed bacteria is calculated by the following equation: ((average CFU with gentamicin)/(average CFU without gentamicin))*100. **B.** Gene expression in RAW264.7 macrophages infected with WT or sasD A50.1 MRSA for 3 hours. **C-E.** Bone marrow-derived macrophages (BMDMs) were infected with WT or sasD A50.1 MRSA for 3 hours at an MOI of 50 in the absence of antibiotics. **C.** Viability measured by trypan blue staining of BMDMs 3 hours post infection. **D-E**. Representative images (**D**) and quantification of western blot analyses (**E**) of BMDM levels of IL-1β and caspase 1. One to three wells were combined per sample and protein levels are normalized to beta actin in each sample. Arrows denote which band was used for quantification. Statistics tested by Mann-Whitney, * p<0.05, *** p<0.001. N=4-7, combination of several experiments, data graphed as mean ± SEM.

## Discussion

The majority of the viral super-infection literature focuses on the differences in immune responses between bacterial pneumonia and influenza bacterial super-infection. It is well documented that preceding influenza greatly impairs the antibacterial response within the lung(12, 13). Few studies have examined bacterial factors in single or super-infection in the lung. The literature suggests that increased inflammation and tissue damage lead to increased adhesion within the lung, contributing to increases in bacterial burden(9, 10). However, to our knowledge, there has been no specific testing of bacterial adhesion components *in vivo* during single or viral super-infection in the lung. Most studies that have investigated *S. aureus* virulence factors in the lung have focused on secreted toxins, such as the alpha toxin(19, 27–30). While toxin-mediated damage contributes to lung pathology, the alpha toxin has been shown to decrease adhesion to lung epithelial cells(31). Thus, we wanted to determine if proteins with known adhesion properties influenced the outcomes of single or super-infection.

Our data supports the finding that changes due to influenza infection are the primary driver of super-infection, with influenza increasing bacterial burden, immune recruitment, and acute lung injury seen in the model. Interestingly, regardless of what CWA was removed, influenza appeared to “level the playing field” for the mutants, with endpoints being much higher and tighter grouped in super-infection than in bacterial pneumonia alone. *S. aureus* strains seen in super-infected individuals are less virulent and more closely related to nasal colonizing strains than those strains found in bacterial pneumonia patients(32). This is likely due to the increased inflammation and damage within lung as well as a more dysregulated immune response during super-infection leading to less aggressive colonizing strains taking hold in the lung. However, viral-bacterial synergism is likely adding to this phenomenon, as influenza can increase both internalization and adhesion of bacteria within the lung(33, 34). This is not specific to influenza, as the same phenomenon is seen in rhinovirus-*S. aureus* super-infections(1, 35).

We saw more variability in the endpoints studied during bacterial pneumonia, likely because adhesion in the lung is more difficult in a homeostatic state. SasG has a known role in biofilm formation(36, 37), which may explain the decrease in burden seen in bacterial pneumonia. SasG has also been shown to adhere to human desquamated nasal epithelial cells via an unknown ligand(38), so it unclear if lacking SasG would have a pronounced impact on murine lung cell adhesion. IsdB is the receptor for hemoglobin and part of the heme acquisition system to attain iron, an important bacterial nutrient(39). While this protein has higher affinity for human hemoglobin, it still plays a role in murine models(39, 40). Thus, it is unsurprising that lacking this CWA had an impact on bacterial survival in both bacterial pneumonia and influenza super-infection. In bacterial pneumonia, *fnbB::Tn* had elevated protein in the BAL, which may be caused by the high bacterial burden and immune cell infiltrate. FnbB, along with FnbA (not tested in this study), have been shown to play a role in invasion into nonprofessional phagocytes via fibronectin-integrin α_5_β_1_ interactions(41–43). This phenomenon has been shown *in vitro* for alveolar epithelium and a FnbB deletion mutant was found to have increased protein leak in a rat model of pneumonia(44). This suggests that internalization of *S. aureus*, and subsequent immune evasion, may reduce inflammation in the lung. SdrD is known to play a role during nasal colonization as well as help promote survival of *S. aureus* in the blood(45–47). Lacking this CWA may allow for reduced number of immune cells during bacterial pneumonia as well as lower cytokine expression.

CWAs are known to bind to several proteins within the host such as fibrinogen and fibronectin(48). In this study we did not explore bacterial adhesion to specific ligands, but it is likely a combination of several ligands, as described at other host sites such as the nose(49). CWAs also have overlapping ligands, such as ClfA, ClfB, FnbA, and FnbB all binding fibrinogen(48). Because we only looked at single CWA mutants, some of the functions of these proteins in bacterial pneumonia and super-infection could be masked.

Even though the CWA mutants had more clear phenotypes in bacterial pneumonia compared to super-infection, the cytokine signature in both settings appears to be driven by the expression of these CWAs. The mutants found in each cluster were consistent in both bacterial pneumonia and super-infection, with the exception of *sasG::Tn*. This suggests that while a majority of the inflammation in the lung is driven by influenza, at least some part of the immune response is shaped by the presence of these CWAs on the cell surface of the bacteria. As SasG has a known role in biofilm formation and influenza is known to induce dissemination of *S. aureus* biofilms(15, 36, 37), this effect could influence how the immune system reacts to this mutant. A majority of the MSCRAMM proteins (*clfA::Tn, clfB::Tn, sdrC::Tn, sdrE::Tn*) cluster together in the high inflammation cluster. This is what we expected to find, as these proteins have similar domains used for ligand binding and this may influence the immune response(22). ClfA has been shown to be a T cell activator driving Th1 and Th17 activation(50). While we did not see any significant changes in IL-2 or IFNγ, we did see a nearly significant decrease in IL-17A (p=0.0571) with the *clfA::Tn* mutant. Unsurprisingly, the Sortase A mutant, which lacks all CWAs on the cell surface, had the lowest expression of cytokines. It is important to note that the Sortase A mutant still makes all the CWAs, but they are secreted into the environment instead of covalently attached to the cell wall. However, it does suggest that the influence on immune signaling is greatest when the CWAs are still attached to the bacteria. However, more testing would be needed in defining the portions of each CWA responsible for altering immune signaling.

During our screen we found that the *sasD::Tn* mutant had decreased levels of myeloid cytokines and increased gene expression of lung homeostatic markers in bacterial pneumonia. We found similar findings with a transduced mutant. Most striking was the delay in mortality in mice infected with sasD A50.1, which may be explained by the decrease in bacterial survival seen with sasD A50.1 infection compared to the WT strain. What could be causing this decrease in bacterial survival is unknown, as this protein is uncharacterized. While it is known that SasD has a punctate surface expression versus a ring-like distribution of most CWAs(51, 52), it is unclear if this is contributing to the differences seen in our model. This decrease in bacterial survival of sasD A50.1 could explain the inflammatory differences seen early and late during bacterial pneumonia.

At 6 hours post infection, there is a significant reduction in the macrophages in the lung, as well as IL-1β. It is known that macrophage derived IL-1β can induce excessive inflammation and pathology in the lung(53, 54). The reduction in IL-1β could be explained by the decrease in macrophages early during infection. This decrease in inflammation continued at 24 hours post infection, where there was a reduction in the levels of neutrophils, which can cause excessive damage themselves(55). While we did not see functional changes in neutrohpils via qPCR, we did see a decrease in the neutrophil:eosinophil ratio within the lung at 24 hours post infection with sasD A50.1. Eosinophils have been implicated in antibacterial immunity(56), and the increase ratio of eosinophils could help control the bacterial burden in the lung. It has been shown that IL-33 induction of type 2 responses is protective in lethal models of *S. aureus* sepsis and pneumonia by counterbalancing pro-inflammatory responses(57, 58). While we did not see any differences in IL-33 (data not shown) or gross pathology at 24 hours post infection, we did see a reduction in type 17 cytokines and neutrophils, which has been shown to be protective in patients with *S. aureus* infection(58, 59). Thus, the reduction in inflammation or alteration of inflammatory cell ratios could help explain the delayed mortality seen in mice.

Since we saw a change in IL-1β production both early and late during infection, we decided to examine the inflammasome. *S. aureus* is known to prime and activate the NLRP3 inflammasome via pore-forming toxins, such as the alpha toxin(26). The NLRP3 inflammasome activates caspase 1, which cleaves pro-IL-1β(30). We did see a significant downregulation of *il-1β* and *nlrp3* transcripts but not the more common ASC (*pycard*) component, suggesting that potentially the priming step of the NLRP3 inflammasome expression may be reduced. Priming of the NLRP3 inflammasome is thought to be due to sensing of *S. aureus* lipoproteins and toll-like receptor (TLR) 2 and 4 signaling(26, 60). While we did not see changes in expression in TLR-2 or −4 in macrophages (data not shown), we cannot rule out the possibility that SasD may be involved in the sensing of *S. aureus*. When infected with sasD A50.1, RAW264.7 cells had a reduction in *il-1β* and *tnfa* without a significant change in bacterial burden or bacterial phagocytosis. In BMDMs, we saw increased viability when infected with sasD A50.1 compared to WT *S. aureus*. While we did not see any differences in pro-IL-1β, caspase 1, or caspase 1 cleavage at 3 hours post infection, there may be other cell death pathways involved such as necroptosis. Blocking necroptosis has been shown to reduce bacterial burden and damage during *S. aureus* pneumonia(29), similar to the sasD A50.1 *in vivo* phenotype, and can drive the NLRP3 inflammasome and pyroptosis induction(61). Thus, the decrease in IL-1β could be due to changes in cell death pathways that result into the NLRP3 inflammasome activation in macrophages.

In conclusion, we identified a critical role for SasD in bacterial pneumonia associated with increased bacterial burden, inflammation, and mortality. SasD may contribute to survival of *S. aureus* in the lung as there was decreased bacterial survival in the mutant at both 6 and 24 hours post infection. SasD promotes induction of early IL-1β production in macrophages, which consequentially recruits neuctrophils into the lung at later timepoints, leading to increased inflammation. These data suggest that early targeting of SasD in the lung may reduce future inflammation signaling during staphylococcal pneumonia.

## Methods

### Mice

Six- to eight-week-old male and female WT C57BL/6NTac mice were purchased from Taconic Farms. Mice were maintained under pathogen-free conditions within the animal facilities at the UPMC Children’s Hospital of Pittsburgh. All studies were performed on sex- and age-matched mice. Animal studies were conducted with approval from the University of Pittsburgh Institutional Animal Care and Use Committee.

### *S. aureus* strains

USA300 MRSA strain JE2(62) was the WT strain for all studies. All strains used in study are listed in Table 1 and are derived from the Nebraska Transposon Mutant Library(62) (BEI Resources), with strains *srtA::Tn, sdrE::Tn, sdrC::Tn, fnbB::Tn, isdB::Tn, sasG::Tn*, and *sasD::Tn* gifted from Dr. Ken Urish, University of Pittsburgh. All mutants were confirmed by PCR using gene-and transposon-specific primers in Table 2. Strain sasD A50.1 was generated via phage 11 transduction of *sasD::Tn* lysate into the wildtype JE2 strain, selected with 5 μg/ml erythromycin and confirmed by PCR (Table 2). *S. aureus* strains were grown in Tryptic Soy Broth (BD Bacto™) overnight at 37° C at 250 rpm. Overnight cultures were diluted 1:100 and grown until OD_660_~1, approximating logarithmic growth phase. MRSA dose was calculated using OD660 measurement of the culture and application of a calculated extinction coefficient. For growth curves, overnight cultures were diluted 1:200 in a 96-well plate in sexaplicate. Plates were grown at 37°C at 282 rpm in a Synergy H1 Hybrid Multi-Mode Reader (BioTek). Optical density measurements at 660 nm were taken every 30 minutes. Growth rate (μ) was calculated from at least two independent experiments using the equation A_t_=A_t-1_*e^μt^. The μ_max_ was calculated as the average of the three highest μ rates.

**Table 1:** *S. aureus* strains used in this study

| Strain | Description | NTML # | Reference |
| --- | --- | --- | --- |
| JE2 | NTML Wildtype strain |  | (62) |
| JE2 *srtA::Tn* | NTML mutant | NE1787 | (62) |
| JE2 *clfA::Tn* | NTML mutant | NE543 | (62) |
| JE2 *clfB::Tn* | NTML mutant | NE391 | (62) |
| JE2 *fnbB::Tn* | NTML mutant | NE728 | (62) |
| JE2 *sdrC::Tn* | NTML mutant | NE432 | (62) |
| JE2 *sdrD::Tn* | NTML mutant | NE1289 | (62) |
| JE2 *sdrE::Tn* | NTML mutant | NE98 | (62) |
| JE2 *isdB::Tn* | NTML mutant | NE1102 | (62) |
| JE2 *sasG::Tn* | NTML mutant | NE825 | (62) |
| JE2 *sasD::Tn* | NTML mutant | NE1032 | (62) |
| sasD A50.1 | Transduced strain from JE2 *sasD::Tn* |  | This study |

**Table 2:**
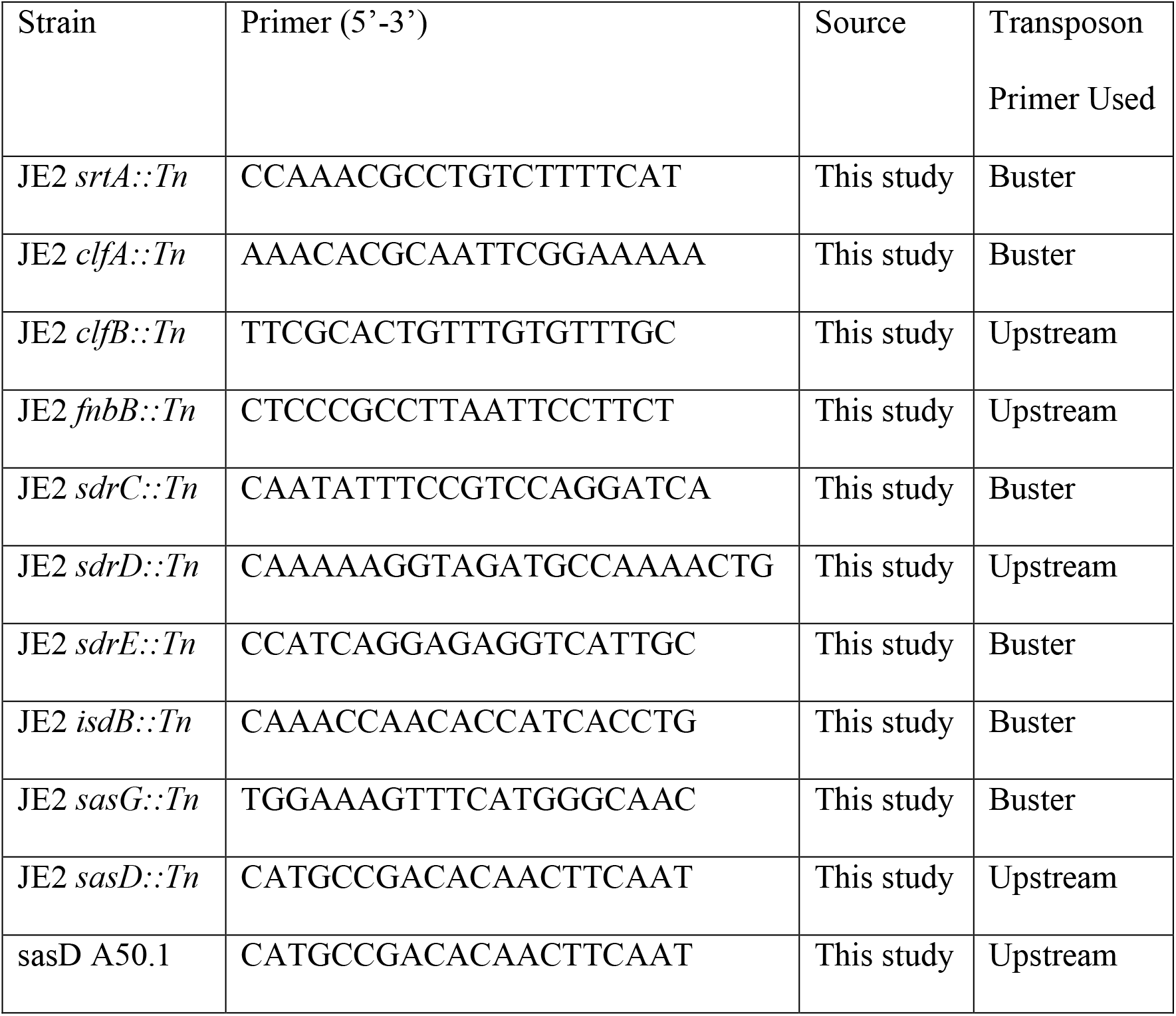
*S. aureus* primers used

### Murine Models

Influenza A/PR/8/34 (H1N1) was grown in chicken eggs as previously described(63). Mice were inoculated with PBS vehicle or 100 plaque forming units (PFU) of influenza in 50 μl of sterile PBS. Six days later, mice were infected with 1×10 colony forming units (CFU) of MRSA in 50 μl of sterile PBS. All infections were performed via oropharyngeal aspiration. Mice were harvested 6 or 24 hours after MRSA challenge using pentobarbital injection (300 mg/kg) and cervical dislocation. In mortality studies, a dose of 2×10 CFU was used. During harvest, the lung was lavaged with 1 ml sterile PBS. BAL cells were pelleted and red blood cells were lysed (ACK lysis buffer, Gibco). Cells were resuspended, placed on slides via cytospin, stained with Hema 3 (Thermo Fisher), and quantified. The right upper lung lobe was homogenized in 1 ml PBS and plated on tryptic soy agar for determination of bacterial burden. The remaining right lung was frozen in liquid nitrogen and stored at −80°C for gene expression analysis. The left lobe was perfused with 10% formalin and embedded in paraffin. Lung sections were stained with hematoxylin and eosin and inflammatory features were evaluated via microscopy after sample blinding. For competitive index studies, mice were inoculated with a 1:1 ratio of JE2 and sasD A50.1 strains at a total of 1×10 CFU. Whole lungs were homogenized in 2 ml of sterile PBS and plated on tryptic soy agar with and without erythromycin (5 μg/ml). Competitive index was calculated as the ratio of sasD A50.1:JE2 CFU at sacrifice divided by the ratio at the time of innoculation.

### Macrophage experiments

RAW264.7 cells were used and BMDMs were isolated as previously described(64). For experiments, 7×10^5^ cells were plated in 6-well plates, infected at an MOI 10 (RAW264.7) or 50 (BMDMs) and spun at 250 xg for 5 minutes at 4° C to synchronize infection. For RAW264.7 experiments, cells were infected for 1 hour in the absence of antibiotics, media was replaced with antibiotic- and serum-free media with and without gentamicin (100 ug/ml) for 1 hour, then replaced with antibiotic-free media for an additional hour. At collection, cells were lysed with 1% Triton X-100 at room temperature for 10 minutes and 50 μl was collected for CFU determination. Phagocytosis was calculated by the equation ((CFU+gentamicin)/(CFU-gentamicin))*100. RLT (Qiagen) was added to the wells and collected and ran through a Qiashredder and frozen at −80°C until RNA extraction. For BMDM experiments, cells were rested overnight, treated with 10 ng/ml IFNγ (R&D Systems) for 24 hours. BMDMs were infected for 3 hours, washed and resuspended in antibiotic free RPMI media. BMDM viability was determined by trypan blue (Gibco) staining and the Countess 3 automatic cell counter (Invitrogen). BMDMs wells were combined and incubated in RIPA buffer (25mM Tris, 150 mM NaCl, 1% NP-40, 0.1% SDS, 5 mM EDTA, 0.5% sodium deoxycholate) for 30 minutes at 4°C with agitation, centrifuged at 10,000 rpm for 10 minutes at 4°C, and frozen at −80°C until Western Blot. Primary antibodies were rabbit anti-IL-1β (Abcam 254360), rabbit anti-caspase 1 (Abcam 138483), rabbit anti-caspase p20 (Invitrogen PA5-99390), and mouse anti-β-actin (Cell Signaling 8H10D10). Samples were thawed, proteins were quantified using BCA protein assay (Pierce), boiled in Laemmli buffer (Bio-Rad), and loaded on a 4-20% gel (Bio-Rad). Proteins were transferred to a PVDF membrane using the Trans-Blot Turbo transfer system (Bio-Rad). Blots were probed with primary antibodies and donkey anti-mouse or goat anti-rabbit secondary antibodies conjugated to IRDye^®^ 800CW or 680RD flurophores (LI-COR). Blots were imaged using the Odyssey CLx and analyzed using Image Studio (LI-COR). Relative protein expression is normalized to beta-actin levels in each sample.

### RNA extraction and qPCR

RNA was extracted from mouse lungs using the Absolutely Total RNA Purification Kit (Agilent). RNA extraction from cell culture experiments were performed using the Qiagen RNeasy kit (Qiagen). RNA was quantified and converted to cDNA using iScript™ cDNA Synthesis Kit (Bio-Rad). Quantitative PCR was performed using SsoAdvanced Universal Probes Supermix (Bio-Rad) and TaqMan primer-probe sets (ThermoFisher Scientific) listed in Table 3 on the CFX96 Touch Real-Time PCR Detection System (Bio-Rad). Gene expression was calculated using the ΔΔct method using *hprt* as a housekeeping gene and normalized to the average WT *S. aureus* values unless otherwise stated.

**Table 3:** TaqMan Primer-Probes used in study

| Gene | Catalog Number |
| --- | --- |
| IL-17a | Mm00439618_m1 |
| IL-23a | Mm00518984_m1 |
| Cxcl1 | Mm04207460_m1 |
| IL-1β | Mm00434228_m1 |
| NLRP3 | Mm00840904_m1 |
| Pycard | Mm00445747_g1 |
| Foxj1 | Mm01267279_m1 |
| TJP1 | Mm00493699_m1 |
| Cav1 | Mm00483057_m1 |
| Sftpc | Mm00488144_m1 |
| Scgb1a1 | Mm00442046_m1 |
| Muc5b | Mm00466376_m1 |
| MPO | Mm01298424_m1 |
| ELANE | Mm00469310_m1 |
| CTSG | Mm00456011_m1 |

### Multiplex and Heatmap analysis

Lung homogenate cytokines were assessed using the Bio-Plex Pro Mouse Cytokine 23-plex Assay (Bio-Rad). For clustering analysis, all data was combined and samples with missing data and MIP-1α, due to poor detection, were excluded. The average for each cytokine per mutant was used. Using R (version 4.1.0) in RStudio (version 1.4.1717), data was log-transformed and scaled to Z score and clustered by cytokine using the hclust function and Pearson correlation. The resulting heatmap was visualized using Heatmap.2 function in the gplots package.

### Statistical Analysis

Data were analyzed using Prism 8 (GraphPad). Analyses comparing two groups were performed using Mann-Whitney test or an unpaired t test. For analyses assessing more than two groups, Kruskall-Wallis with Dunn’s multiple comparisons correction was used. Analyses comparing two variables were tested via Two-way analysis of variance (ANOVA) with Sidak’s multiple comparisons correction. Mortality data were analyzed by a log rank (Mantel-Cox) test. All figures show combined data from multiple replicate studies and are graphed as mean ± standard error of the mean (SEM). N values are numbers of animals per independent experiment. Statistical significance (p ≤ 0.05) is indicated in figure legends, with p values between 0.05 and 0.1 displayed numerically.

## Supplemental Figure Legends

**Supplemental Figure 1:**
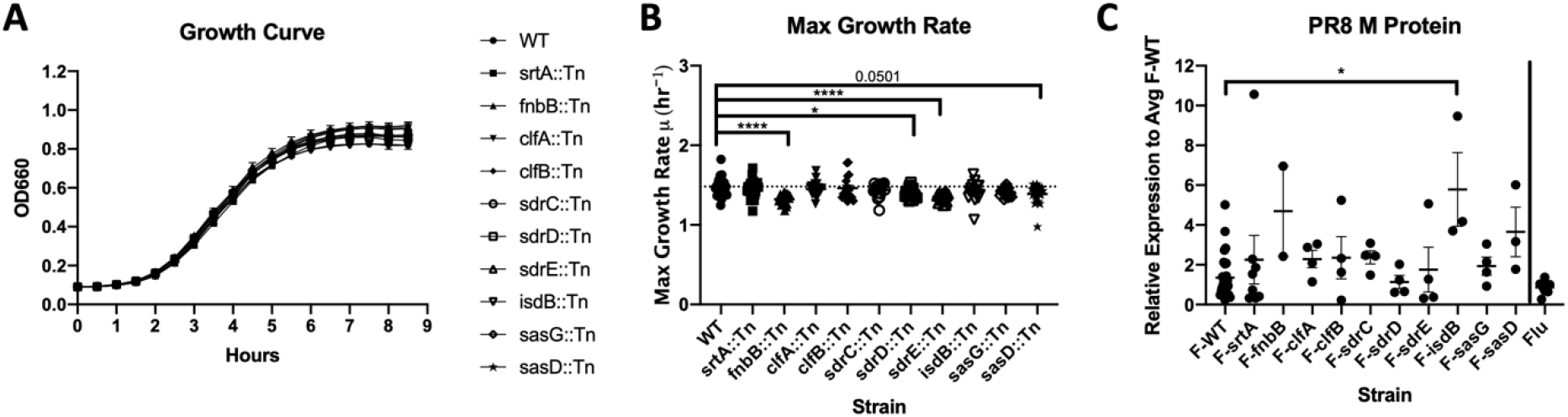
Bacterial Growth and Viral Burden. **A.** WT or mutant MRSA (see graphs) were grown overnight in tryptic soy broth and diluted 1:200 in a 96-well microtiter plate in sexaplicate. Plates were grown at 37°C at 282 rpm continuously. Measurements at 660 nm were taken every 30 minutes. Combination of at least 2 experiments per strain. **B.** Max growth rate of WT or mutant MRSA strains (see graphs). Growth rate (μ) was calculated off at least two independent experiments using the equation A_t_=A_t-1_*e^μt^ (see methods). The μ_max_ was calculated as the average of the three highest μ rates. **C.** Relative expression of influenza PR8 M protein via qPCR normalized to the average F-WT values. Statistics tested by Kruskal-Wallis with Dunn’s multiple comparisons correction (B-C). * p<0.05, ****p<0.0001. N=2-6, combination of several experiments, data graphed as mean ± SEM. srtA: Sortase A, fnbB: fibronectin binding protein B, clfA/B: clumping factor A/B, sdrC/D/E: serine-aspartate repeat containing protein C/D/E, isdB: iron-regulated surface determinant B, sasD/G: *S. aureus* surface protein D/G.

**Supplemental Figure 2:**
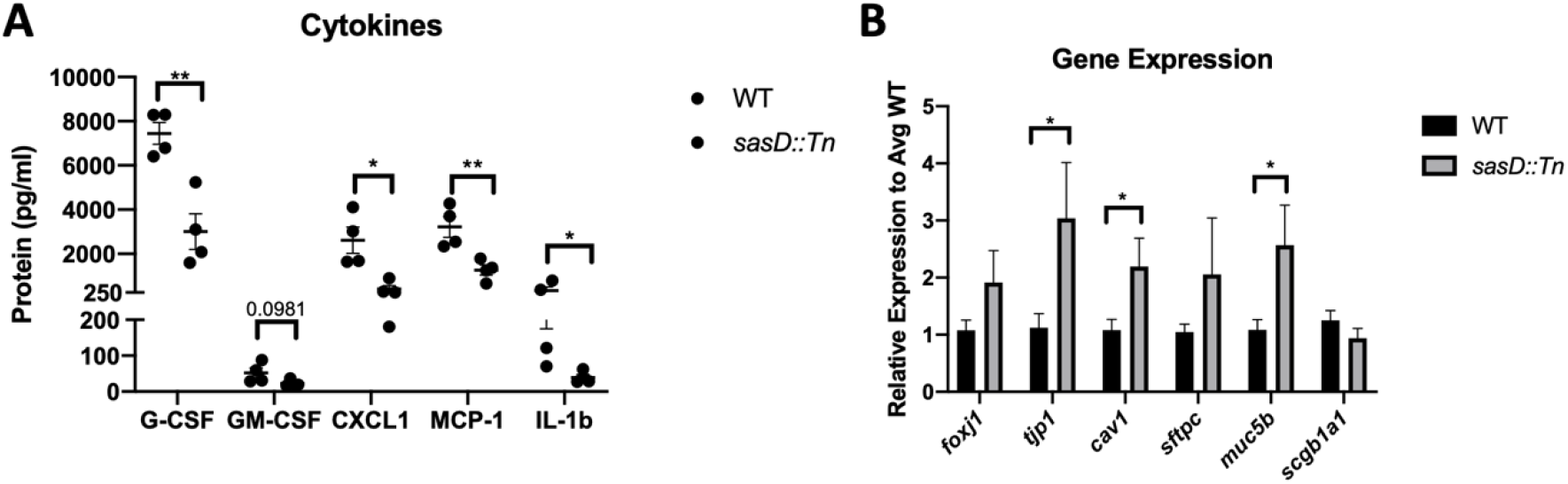
SasD Increases Inflammatory Cytokine and Decreases Lung Homoestatic Gene Expression. **A.** Lung homogenate protein levels of cytokines in mice infected with WT or *sasD::Tn* MRSA during bacterial pneumonia. **B.** Gene expression of lung epithelial markers in mice infected with WT or *sasD::Tn* MRSA during bacterial pneumonia. Statistics tested by unpaired t test. * p<0.05, **p<0.01. N=2, combination of several experiments, data graphed as mean ± SEM.

**Supplemental Figure 3:**
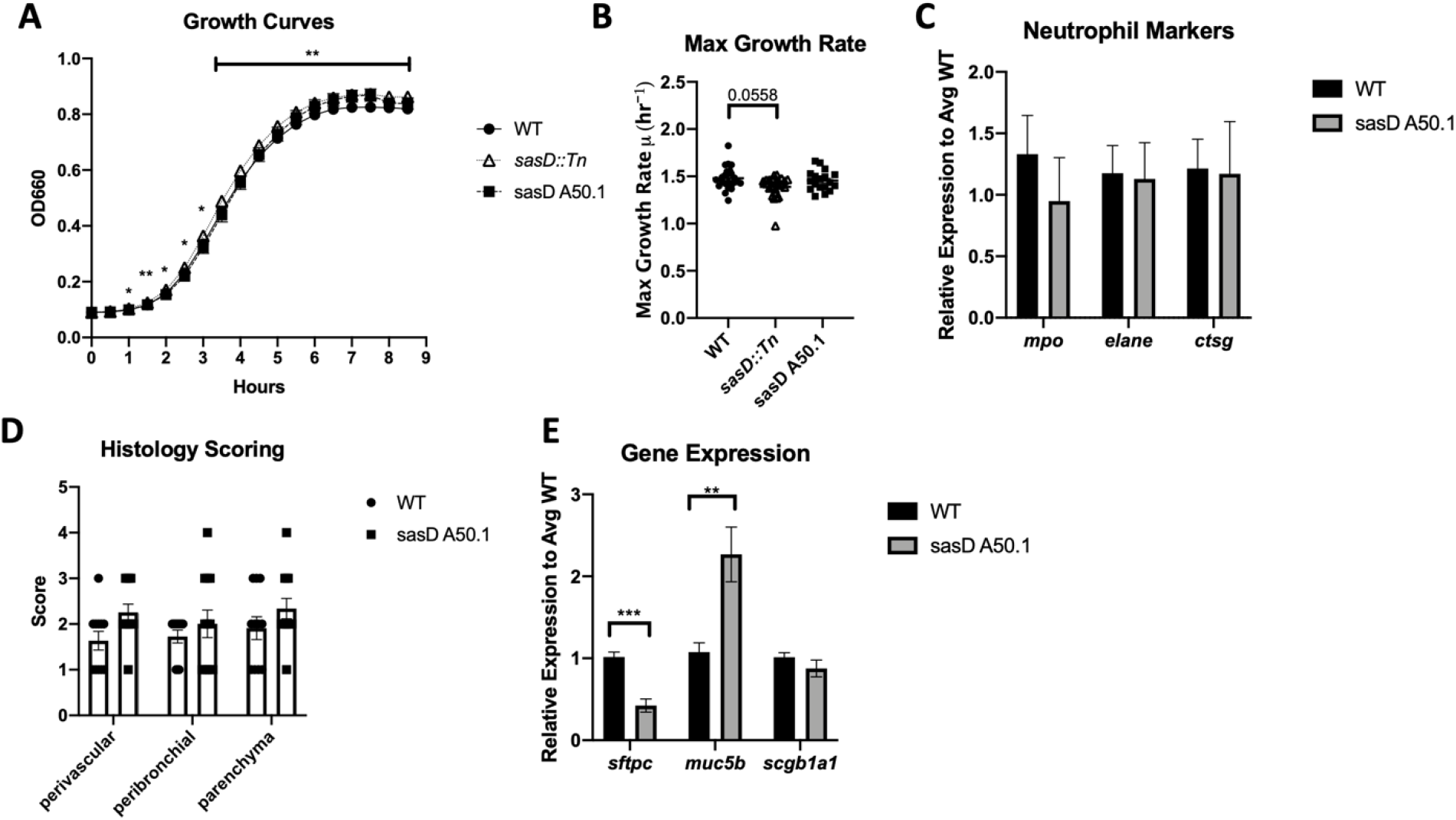
Characterization of SasD during Pneumonia. **A.** WT or mutant MRSA (see graphs) were grown overnight in tryptic soy broth and diluted 1:200 in a 96-well microtiter plate in sexaplicate. Plates were grown at 37°C at 282 rpm continuously. Measurements at 660 nm were taken every 30 minutes. Combination of at least 2 experiments per strain. Statistical significance is between WT and *sasD::Tn* MRSA strains. **B.** Max growth rate of WT or mutant MRSA strains (see graphs). Growth rate (μ) was calculated off at least two independent experiments using the equation A_t_=A_t-1_*e^μt^ (see methods). The μ_max_ was calculated as the average of the three highest μ rates. **C.** Gene expression of neutrophil markers relative to the average WT values. **D.** Histology scoring of H&E-stained lung sections. **E.** Gene expression of lung epithelial markers relative to the average WT values. Statistics done by Mixed-effects model with Dunnett’s multiple comparisons correction (A), Kruskal-Wallis Test with Dunn’s multiple comparisons correction (B), unpaired T test (C-E). * p<0.05, ** p<0.01, *** p<0.001, N=2-6, combination of multiple experiments, data graphed as mean ± SEM.

**Supplemental Figure 4:**
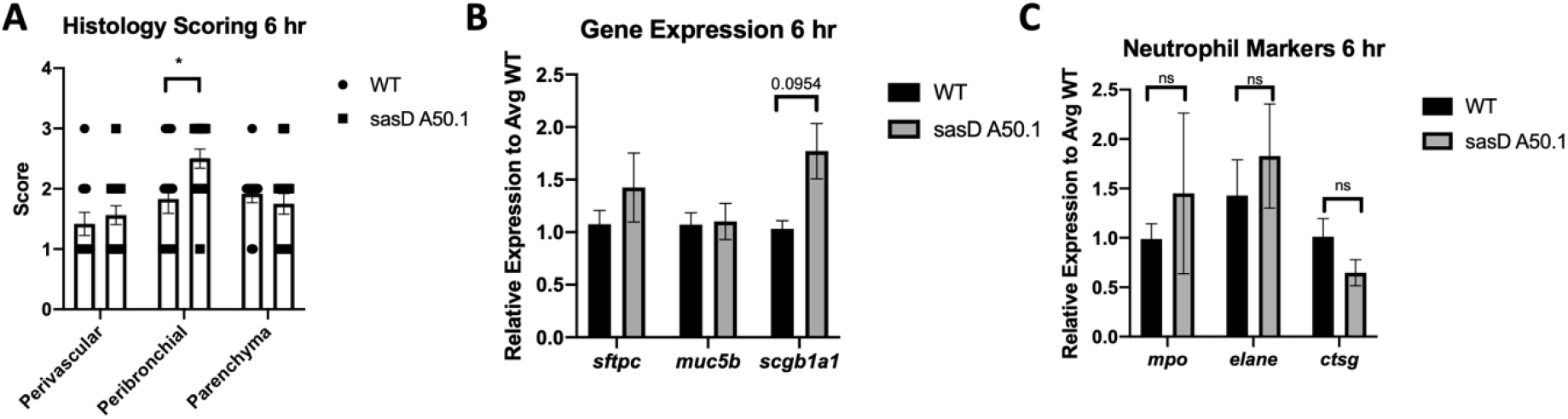
Characterization of early SasD infection. **A.** Histology scoring of H&E-stained lung sections. **B-C.** Gene expression of lung epithelial (**B**) and neutrophil (**C**) markers relative to the average WT values. Statistics done by Two-way ANOVA with Sidak’s multiple comparisons correction (A), unpaired T test (B-C). * p<0.05, n=4-6, combination of multiple experiments, data graphed as mean ± SEM.

## References

1. Morgene MF, Botelho-Nevers E, Grattard F, Pillet S, Berthelot P, Pozzetto B, Verhoeven PO. 2018. Staphylococcus aureus colonization and non-influenza respiratory viruses: Interactions and synergism mechanisms. Virulence 9:1354–1363.

2. Adalbert JR, Varshney K, Tobin R, Pajaro R. 2021. Clinical outcomes in patients co-infected with COVID-19 and Staphylococcus aureus: a scoping review. BMC Infect Dis 21:985.

3. Kourtis AP, Hatfield K, Baggs J, Mu Y, See I, Epson E, Nadle J, Kainer MA, Dumyati G, Petit S, Ray SM, Ham D, Capers C, Ewing H, Coffin N, McDonald LC, Jernigan J, Cardo D, group EIPMa. 2019. Vital Signs: Epidemiology and Recent Trends in Methicillin-Resistant and in Methicillin-Susceptible Staphylococcus aureus Bloodstream Infections - United States. MMWR Morb Mortal Wkly Rep 68:214–219.

4. Paget J, Spreeuwenberg P, Charu V, Taylor RJ, Iuliano AD, Bresee J, Simonsen L, Viboud C, Teams* GSI-aMCNaGC. 2019. Global mortality associated with seasonal influenza epidemics: New burden estimates and predictors from the GLaMOR Project. J Glob Health 9:020421.

5. Rice TW, Rubinson L, Uyeki TM, Vaughn FL, John BB, Miller RR, Higgs E, Randolph AG, Smoot BE, Thompson BT, Network NA. 2012. Critical illness from 2009 pandemic influenza A virus and bacterial coinfection in the United States. Crit Care Med 40:1487–98.

6. (CDC) CfDCaP. 2012. Severe coinfection with seasonal influenza A (H3N2) virus and Staphylococcus aureus--Maryland, February-March 2012. MMWR Morb Mortal Wkly Rep 61:289–91.

7. Shah NS, Greenberg JA, McNulty MC, Gregg KS, Riddell J, Mangino JE, Weber DM, Hebert CL, Marzec NS, Barron MA, Chaparro-Rojas F, Restrepo A, Hemmige V, Prasidthrathsint K, Cobb S, Herwaldt L, Raabe V, Cannavino CR, Hines AG, Bares SH, Antiporta PB, Scardina T, Patel U, Reid G, Mohazabnia P, Kachhdiya S, Le BM, Park CJ, Ostrowsky B, Robicsek A, Smith BA, Schied J, Bhatti MM, Mayer S, Sikka M, Murphy-Aguilu I, Patwari P, Abeles SR, Torriani FJ, Abbas Z, Toya S, Doktor K, Chakrabarti A, Doblecki-Lewis S, Looney DJ, David MZ. 2016. Bacterial and viral co-infections complicating severe influenza: Incidence and impact among 507 U.S. patients, 2013-14. J Clin Virol 80:12–9.

8. Randolph AG, Vaughn F, Sullivan R, Rubinson L, Thompson BT, Yoon G, Smoot E, Rice TW, Loftis LL, Helfaer M, Doctor A, Paden M, Flori H, Babbitt C, Graciano AL, Gedeit R, Sanders RC, Giuliano JS, Zimmerman J, Uyeki TM, Pediatric Acute Lung Injury and Sepsis Investigator’s Network and the National Heart Ln, and Blood Institute ARDS Clinical Trials Network. 2011. Critically ill children during the 2009-2010 influenza pandemic in the United States. Pediatrics 128:e1450–8.

9. McCullers JA. 2014. The co-pathogenesis of influenza viruses with bacteria in the lung. Nat Rev Microbiol 12:252–62.

10. Siemens N, Oehmcke-Hecht S, Mettenleiter TC, Kreikemeyer B, Valentin-Weigand P, Hammerschmidt S. 2017. Port d’Entrée for Respiratory Infections - Does the Influenza A Virus Pave the Way for Bacteria? Front Microbiol 8:2602.

11. Shirey KA, Perkins DJ, Lai W, Zhang W, Fernando LR, Gusovsky F, Blanco JCG, Vogel SN. 2019. Influenza “Trains” the Host for Enhanced Susceptibility to Secondary Bacterial Infection. mBio 10.

12. Metzger DW, Sun K. 2013. Immune dysfunction and bacterial coinfections following influenza. J Immunol 191:2047–52.

13. Robinson KM, Kolls JK, Alcorn JF. 2015. The immunology of influenza virus-associated bacterial pneumonia. Curr Opin Immunol 34:59–67.

14. Rynda-Apple A, Robinson KM, Alcorn JF. 2015. Influenza and Bacterial Superinfection: Illuminating the Immunologic Mechanisms of Disease. Infect Immun 83:3764–70.

15. Reddinger RM, Luke-Marshall NR, Hakansson AP, Campagnari AA. 2016. Host Physiologic Changes Induced by Influenza A Virus Lead to Staphylococcus aureus Biofilm Dispersion and Transition from Asymptomatic Colonization to Invasive Disease. mBio 7.

16. Wang C, Armstrong SM, Sugiyama MG, Tabuchi A, Krauszman A, Kuebler WM, Mullen B, Advani S, Advani A, Lee WL. 2015. Influenza-Induced Priming and Leak of Human Lung Microvascular Endothelium upon Exposure to Staphylococcus aureus. Am J Respir Cell Mol Biol 53:459–70.

17. Puchelle E, Zahm JM, Tournier JM, Coraux C. 2006. Airway epithelial repair, regeneration, and remodeling after injury in chronic obstructive pulmonary disease. Proc Am Thorac Soc 3:726–33.

18. McCullers JA, Bartmess KC. 2003. Role of neuraminidase in lethal synergism between influenza virus and Streptococcus pneumoniae. J Infect Dis 187:1000–9.

19. Hook JL, Islam MN, Parker D, Prince AS, Bhattacharya S, Bhattacharya J. 2018. Disruption of staphylococcal aggregation protects against lethal lung injury. J Clin Invest 128:1074–1086.

20. Foster TJ, Geoghegan JA, Ganesh VK, Höök M. 2014. Adhesion, invasion and evasion: the many functions of the surface proteins of Staphylococcus aureus. Nat Rev Microbiol 12:49–62.

21. Foster TJ. 2019. Surface Proteins of. Microbiol Spectr 7.

22. Foster TJ. 2019. The MSCRAMM Family of Cell-Wall-Anchored Surface Proteins of Gram-Positive Cocci. Trends Microbiol 27:927–941.

23. Schneewind O, Missiakas D. 2014. Sec-secretion and sortase-mediated anchoring of proteins in Gram-positive bacteria. Biochim Biophys Acta 1843:1687–97.

24. Schneewind O, Missiakas D. 2019. Sortases, Surface Proteins, and Their Roles in. Microbiol Spectr 7.

25. Mulcahy ME, Geoghegan JA, Monk IR, O’Keeffe KM, Walsh EJ, Foster TJ, McLoughlin RM. 2012. Nasal colonisation by Staphylococcus aureus depends upon clumping factor B binding to the squamous epithelial cell envelope protein loricrin. PLoS Pathog 8:e1003092.

26. Muñoz-Planillo R, Franchi L, Miller LS, Núñez G. 2009. A critical role for hemolysins and bacterial lipoproteins in Staphylococcus aureus-induced activation of the Nlrp3 inflammasome. J Immunol 183:3942–8.

27. Cohen TS, Hilliard JJ, Jones-Nelson O, Keller AE, O’Day T, Tkaczyk C, DiGiandomenico A, Hamilton M, Pelletier M, Wang Q, Diep BA, Le VT, Cheng L, Suzich J, Stover CK, Sellman BR. 2016. Staphylococcus aureus α toxin potentiates opportunistic bacterial lung infections. Sci Transl Med 8:329ra31.

28. Becker KA, Fahsel B, Kemper H, Mayeres J, Li C, Wilker B, Keitsch S, Soddemann M, Sehl C, Kohnen M, Edwards MJ, Grassmé H, Caldwell CC, Seitz A, Fraunholz M, Gulbins E. 2018. Staphylococcus aureus Alpha-Toxin Disrupts Endothelial-Cell Tight Junctions via Acid Sphingomyelinase and Ceramide. Infect Immun 86.

29. Kitur K, Parker D, Nieto P, Ahn DS, Cohen TS, Chung S, Wachtel S, Bueno S, Prince A. 2015. Toxin-induced necroptosis is a major mechanism of Staphylococcus aureus lung damage. PLoS Pathog 11:e1004820.

30. Grousd JA, Rich HE, Alcorn JF. 2019. Host-Pathogen Interactions in Gram-Positive Bacterial Pneumonia. Clin Microbiol Rev 32.

31. Liang X, Ji Y. 2006. Alpha-toxin interferes with integrin-mediated adhesion and internalization of Staphylococcus aureus by epithelial cells. Cell Microbiol 8:1656–68.

32. Deinhardt-Emmer S, Haupt KF, Garcia-Moreno M, Geraci J, Forstner C, Pletz M, Ehrhardt C, Löffler B. 2019. Pneumonia: Preceding Influenza Infection Paves the Way for Low-Virulent Strains. Toxins (Basel) 11.

33. Rowe HM, Meliopoulos VA, Iverson A, Bomme P, Schultz-Cherry S, Rosch JW. 2019. Direct interactions with influenza promote bacterial adherence during respiratory infections. Nat Microbiol 4:1328–1336.

34. Passariello C, Nencioni L, Sgarbanti R, Ranieri D, Torrisi MR, Ripa S, Garaci E, Palamara AT. 2011. Viral hemagglutinin is involved in promoting the internalisation of Staphylococcus aureus into human pneumocytes during influenza A H1N1 virus infection. Int J Med Microbiol 301:97–104.

35. Passariello C, Schippa S, Conti C, Russo P, Poggiali F, Garaci E, Palamara AT. 2006. Rhinoviruses promote internalisation of Staphylococcus aureus into non-fully permissive cultured pneumocytes. Microbes Infect 8:758–66.

36. Corrigan RM, Rigby D, Handley P, Foster TJ. 2007. The role of Staphylococcus aureus surface protein SasG in adherence and biofilm formation. Microbiology (Reading) 153:2435–2446.

37. Geoghegan JA, Corrigan RM, Gruszka DT, Speziale P, O’Gara JP, Potts JR, Foster TJ. 2010. Role of surface protein SasG in biofilm formation by Staphylococcus aureus. J Bacteriol 192:5663–73.

38. Roche FM, Meehan M, Foster TJ. 2003. The Staphylococcus aureus surface protein SasG and its homologues promote bacterial adherence to human desquamated nasal epithelial cells. Microbiology (Reading) 149:2759–2767.

39. Torres VJ, Pishchany G, Humayun M, Schneewind O, Skaar EP. 2006. Staphylococcus aureus IsdB is a hemoglobin receptor required for heme iron utilization. J Bacteriol 188:8421–9.

40. Cheng AG, Kim HK, Burts ML, Krausz T, Schneewind O, Missiakas DM. 2009. Genetic requirements for Staphylococcus aureus abscess formation and persistence in host tissues. FASEB J 23:3393–404.

41. Dziewanowska K, Patti JM, Deobald CF, Bayles KW, Trumble WR, Bohach GA. 1999. Fibronectin binding protein and host cell tyrosine kinase are required for internalization of Staphylococcus aureus by epithelial cells. Infect Immun 67:4673–8.

42. Menzies BE. 2003. The role of fibronectin binding proteins in the pathogenesis of Staphylococcus aureus infections. Curr Opin Infect Dis 16:225–9.

43. Foster TJ. 2016. The remarkably multifunctional fibronectin binding proteins of Staphylococcus aureus. Eur J Clin Microbiol Infect Dis 35:1923–1931.

44. McElroy MC, Cain DJ, Tyrrell C, Foster TJ, Haslett C. 2002. Increased virulence of a fibronectin-binding protein mutant of Staphylococcus aureus in a rat model of pneumonia. Infect Immun 70:3865–73.

45. Askarian F, Uchiyama S, Valderrama JA, Ajayi C, Sollid JUE, van Sorge NM, Nizet V, van Strijp JAG, Johannessen M. 2017. Serine-Aspartate Repeat Protein D Increases Staphylococcus aureus Virulence and Survival in Blood. Infect Immun 85.

46. Corrigan RM, Miajlovic H, Foster TJ. 2009. Surface proteins that promote adherence of Staphylococcus aureus to human desquamated nasal epithelial cells. BMC Microbiol 9:22.

47. Jenkins A, Diep BA, Mai TT, Vo NH, Warrener P, Suzich J, Stover CK, Sellman BR. 2015. Differential expression and roles of Staphylococcus aureus virulence determinants during colonization and disease. mBio 6:e02272–14.

48. Foster TJ. 2019. Surface Proteins of *Staphylococcus aureus*. Microbiol Spectr 7.

49. Mulcahy ME, McLoughlin RM. 2016. Host-Bacterial Crosstalk Determines Staphylococcus aureus Nasal Colonization. Trends Microbiol 24:872–886.

50. Lacey KA, Leech JM, Lalor SJ, McCormack N, Geoghegan JA, McLoughlin RM. 2017. The Staphylococcus aureus Cell Wall-Anchored Protein Clumping Factor A Is an Important T Cell Antigen. Infect Immun 85.

51. Roche FM, Massey R, Peacock SJ, Day NPJ, Visai L, Speziale P, Lam A, Pallen M, Foster TJ. 2003. Characterization of novel LPXTG-containing proteins of Staphylococcus aureus identified from genome sequences. Microbiology (Reading) 149:643–654.

52. DeDent A, Bae T, Missiakas DM, Schneewind O. 2008. Signal peptides direct surface proteins to two distinct envelope locations of Staphylococcus aureus. EMBO J 27:2656–68.

53. Cohen TS, Prince AS. 2013. Activation of inflammasome signaling mediates pathology of acute P. aeruginosa pneumonia. J Clin Invest 123:1630–7.

54. Pires S, Parker D. 2018. IL-1β activation in response to Staphylococcus aureus lung infection requires inflammasome-dependent and independent mechanisms. Eur J Immunol 48:1707–1716.

55. Lee WL, Downey GP. 2001. Neutrophil activation and acute lung injury. Curr Opin Crit Care 7:1–7.

56. Shamri R, Xenakis JJ, Spencer LA. 2011. Eosinophils in innate immunity: an evolving story. Cell Tissue Res 343:57–83.

57. Krishack PA, Hollinger MK, Kuzel TG, Decker TS, Louviere TJ, Hrusch CL, Sperling AI, Verhoef PA. 2021. IL-33-mediated Eosinophilia Protects against Acute Lung Injury. Am J Respir Cell Mol Biol 64:569–578.

58. Krishack PA, Louviere TJ, Decker TS, Kuzel TG, Greenberg JA, Camacho DF, Hrusch CL, Sperling AI, Verhoef PA. 2019. Protection against Staphylococcus aureus bacteremia-induced mortality depends on ILC2s and eosinophils. JCI Insight 4.

59. Greenberg JA, Hrusch CL, Jaffery MR, David MZ, Daum RS, Hall JB, Kress JP, Sperling AI, Verhoef PA. 2018. Distinct T-helper cell responses to Staphylococcus aureus bacteremia reflect immunologic comorbidities and correlate with mortality. Crit Care 22:107.

60. Wang X, Eagen WJ, Lee JC. 2020. Orchestration of human macrophage NLRP3 inflammasome activation by. Proc Natl Acad Sci U S A 117:3174–3184.

61. Kitur K, Wachtel S, Brown A, Wickersham M, Paulino F, Peñaloza HF, Soong G, Bueno S, Parker D, Prince A. 2016. Necroptosis Promotes Staphylococcus aureus Clearance by Inhibiting Excessive Inflammatory Signaling. Cell Rep 16:2219–2230.

62. Fey PD, Endres JL, Yajjala VK, Widhelm TJ, Boissy RJ, Bose JL, Bayles KW. 2013. A genetic resource for rapid and comprehensive phenotype screening of nonessential Staphylococcus aureus genes. mBio 4:e00537–12.

63. Braciale TJ. 1977. Immunologic recognition of influenza virus-infected cells. I. Generation of a virus-strain specific and a cross-reactive subpopulation of cytotoxic T cells in the response to type A influenza viruses of different subtypes. Cell Immunol 33:423–36.

64. Gopal R, Lee B, McHugh KJ, Rich HE, Ramanan K, Mandalapu S, Clay ME, Seger PJ, Enelow RI, Manni ML, Robinson KM, Rangel-Moreno J, Alcorn JF. 2018. STAT2 Signaling Regulates Macrophage Phenotype During Influenza and Bacterial Super-Infection. Front Immunol 9:2151.

